# OMGene: Mutual improvement of gene models through optimisation of evolutionary conservation

**DOI:** 10.1101/212530

**Authors:** Michael P. Dunne, Steven Kelly

**Affiliations:** Department of Plant Sciences, University of Oxford, South Parks Road, OX1 3RB, UK

## Abstract

**Background:** The accurate determination of the genomic coordinates for a given gene – its *gene model –* is of vital importance to the utility of its annotation, and the accuracy of bioinformatic analyses derived from it. Currently-available methods of computational gene prediction, while on the whole successful, often disagree on the model for a given predicted gene, with some or all of the variant gene models failing to match the biologically observed structure. Many prediction methods can be bolstered by using experimental data such as RNA-seq and mass spectrometry. However, these resources are not always available, and rarely give a comprehensive portrait of an organism’s transcriptome due to temporal and tissue-specific expression profiles.

**Results:** Orthology between genes provides evolutionary evidence to guide the construction of gene models. OMGene (Optimise My Gene) aims to optimise gene models in the absence of experimental data by optimising the derived amino acid alignments for gene models within orthogroups. Using RNA-seq data sets from plants and fungi, considering intron/exon junction representation and exon coverage, and assessing the intra-orthogroup consistency of subcellular localisation predictions, we demonstrate the utility of OMGene for improving gene models in annotated genomes.

**Conclusions:** We show that significant improvements in the accuracy of gene model annotations can be made in both established and *de novo* annotated genomes by leveraging information from multiple species.

## Introduction

The utility of any given genome is dependent on the comprehensiveness and accuracy of its proteome annotation. Inaccuracies in the annotated locations and structures of protein coding genes can lead to myriad downstream errors. These include misinformed conclusions about the biological properties of an organism, as well as errors in transcript quantification, phylogenetic tree inference, protein localisation, and protein structure predictions. It is therefore vital to downstream analysis, both computational and experimental, to ensure that gene annotations are as accurate as possible.

The absolute quantity of publicly available genomic data has grown exponentially over the past two decades, as has the number of taxa represented [1]–[3], owing to the consistently decreasing costs of acquiring whole genome sequences [4], [5]. Accordingly, the feasibility of manual proteome annotation has diminished progressively, with a corresponding increase in reliance on computational gene prediction software. As such there are numerous tools available for the *de novo* and data-assisted prediction of genes [6]. These tools typically rely on genetic signatures such as GC content, codon bias, feature length distributions, and various conserved DNA sequence motifs. Though many of these tools are highly proficient at gene prediction, mistakes are common. Gene prediction tools often disagree on the quantity of genes that they predict [7]–[9]. Furthermore, even when gene predictors agree on the location of a gene, the predicted intron-exon structure for that gene can vary considerably between the different methods [10]. Common such errors include erroneous exon/intron retention/omission, inaccurate exon/intron boundaries, frame errors, misplaced start codons, and fragmentation/fusion of gene models.

When available, the use of extrinsic empirical data, most notably RNA-seq, is the most reliable currently available method for procuring gene models. For example, single contiguous RNA-seq reads obtained from mRNA sequencing can be split across multiple loci when mapped to the genome, providing evidence for the locations of splice junctions. Unfortunately, empirical data is generally not available for all genes in a given species: many genes are expressed in a cell-type or cell-cycle specific manner and for organisms with many disparate tissue types it can be difficult to obtain RNA-seq data that covers the full breadth of the transcriptome [11], [12]. In addition, not all gene sequences are amenable to reliable and accurate alignment, in particular identical duplicate genes and genes that contain repetitive regions found in multiple other genes [13]. Furthermore library preparation protocols and other statistical factors can make reliable inferences difficult [14]– [16]. Finally, there are some aspects of gene models that are simply not revealed by RNA-seq analysis: for example the presence of 5’UTR sequences or internal methionine residues mean that there can often be multiple plausible start codons locations for a given open reading frame (ORF).

Feature locations (splice sites, exons, transcription start sites) have been shown to be highly conserved across evolutionary timescales, often more so than the constituent amino acid sequences they encapsulate [17], [18], despite alternative splicing being a driver of divergence [19]. Given various gene model predictions, it is logical that if multiple highly similar (in sequence and structure) gene models exist for a gene across multiple taxa, they are more likely to be biologically correct than disparate alternatives. By considering *orthogroups* of related genes, one can optimise the similarity of gene models across species by seeking conserved structure across the various taxa. In the absence of extrinsic data, it is parsimonious to choose gene models that maximise intra-orthogroup agreement.

OMGene (Optimise My Gene) aims to improve genome annotations by optimising the agreement between gene models for orthologous genes in multiple species. It is designed to function without the need for additional empirical data, utilising only the local genome sequences for the genes in question, and works on existing predicted gene models. A standalone implementation of the algorithm is available under the GPLv3 licence at https://github.com/mpdunne/omgene. The algorithm is available as a python script, instructions for which, along with example data sets, are included in the git repository.

## Results

### Problem definition, algorithm overview and evaluation criteria

An overview of the OMGene algorithm is provided in Figure 1. OMGene aims to find the most consistent set of representative gene models for a set of inputted genes by seeking to maximise the agreement of their aligned amino acid sequences, returning the single best gene model for each gene. The algorithm constructs gene models based on relatively simple constraints: AUG for start codons; GU or GC for splice donor sites, AG for splice acceptor sites, and UAA, UGA, or UAG for stop codons. Other features such as codon bias or poly-pyrimidine tracts are not considered. OMGene can also use non-canonical translation initiation and splice sites if inputted by the user as a command-line option.

**Figure 1:**
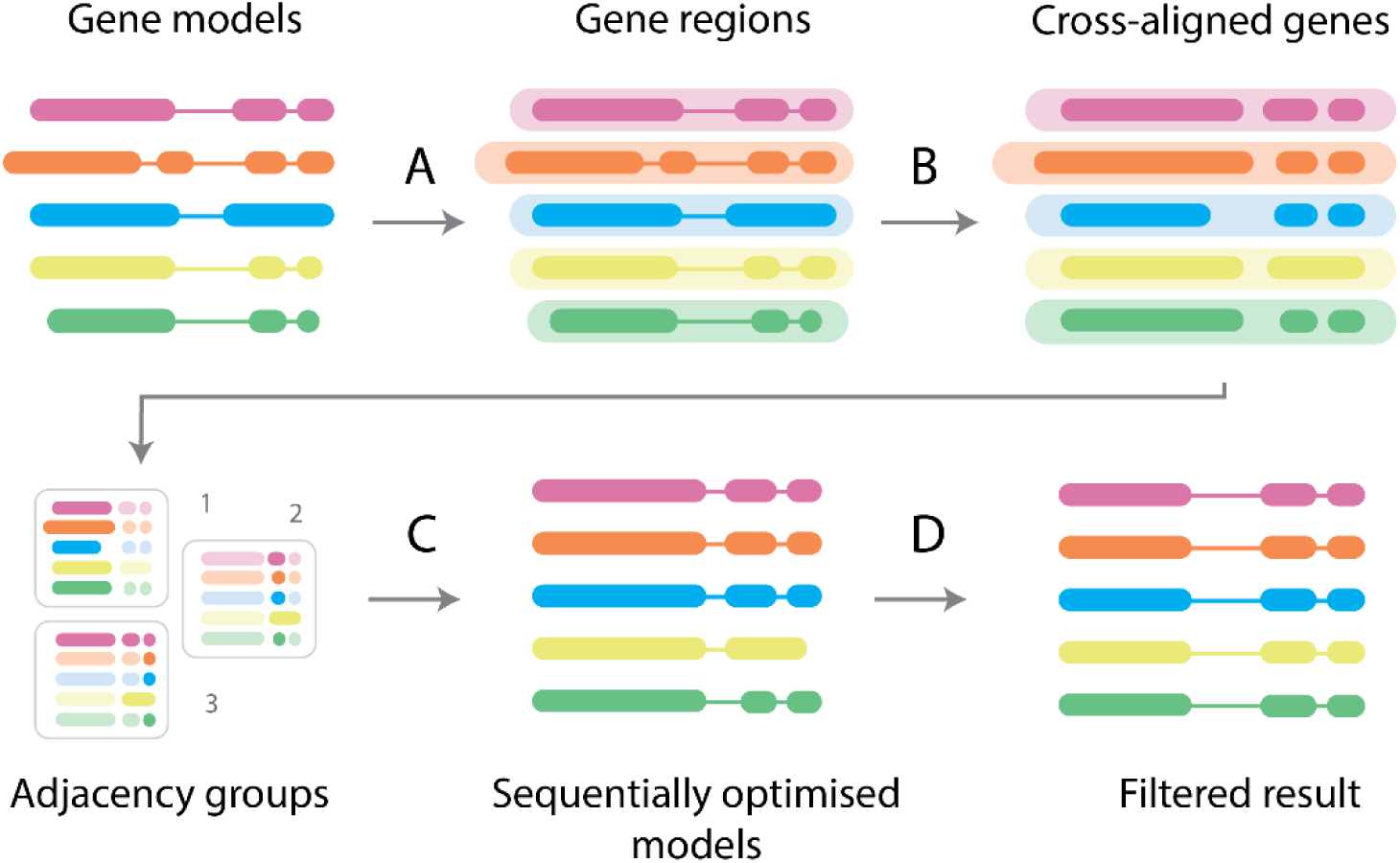
OMGene workflow. Simplified overview of OMGene workflow. A) Gene regions are extracted from around the gene model; B) Exonerate is used to cross-align all constituent exons and full open reading frames to construct basic prototype gene models; C) The exonic regions from these prototype gene models are sorted into adjacency groups, which are then sequentially optimised using the multipartite choice function; D) Results are compared against the original gene models to incorporate potentially overlooked combinations, and filtered under various criteria to produce results.

The input for OMGene is a user-selected set of gene models, in GTF format, which are assumed to belong to a single *orthogroup*. For a given set of species, an *orthogroup* is the set of genes descended from a single ancestral gene in the last common ancestor of those species [20]: these may contain paralogous as well as orthologous genes, though OMGene is principally designed to work on single-copy genes. The suggested pipeline for using OMGene is to determine orthogroups using OrthoFinder [20], and to apply OMGene to a chosen subset of orthogroups.

OMGene uses Exonerate [21] as an initial step to cross-align amino acid sequences from all user-supplied genes to the genomic regions of the genes in question, in order to find conserved translatable features. It then combines this information with the original gene models to produce an initial set of prototype exonic regions, or *gene parts*, for optimisation. The amino acid sequences for these prototype gene models are then aligned, and the constituent gene parts are split into adjacency groups based on overlaps in the alignment (see Methods). Adjacency groups are sequentially appended to the gene models, and the genetic coordinates are recursively adjusted and assessed to optimise the agreement of the amino acid sequences. The resultant gene models are then subject to stringent filtering criteria before the finalised set of gene models are presented as sets of GTF coordinates, amino acid FASTA and CDS FASTA sequences.

To demonstrate the utility of OMGene, it was applied to orthogroups formed from two sets of test species: a set of five fungal species and a set of five plant species (Table 1). OMGene was applied to those orthogroups that contained exactly one gene from each species, referred to as single-copy ubiquitous (SCU) orthogroups. In addition, OMGene was run on the same set but with all genes from two representative species – *A. thaliana* and *S. cerevisiae* – replaced with *de novo* predicted genes, obtained by running the Augustus [22] gene finder on those genomes. These species were chosen as they have the best annotated genomes and thus the existing gene models will provide the best possible training set for Augustus *de novo* prediction. This *de novo* prediction analysis was done to simulate a typical genome-sequencing project where a user has generated a well-trained set of gene models solely using computational prediction.

**Table 1:** Species sets used for algorithm validation

|  | Species Name | Source | Version/Strain | Taxonomy ID | References |
| --- | --- | --- | --- | --- | --- |
| Plant species | *Arabidopsis thaliana* | JGI | TAIR10 | 3702 | [26] |
|  | *Brassica rapa* | JGI | v1.3 | 3711 | [26] |
|  | *Carica papaya* | JGI | ASGPBv0.4 | 3649 | [26] |
|  | *Capsella rubella* | JGI | v1.0 | 81985 | [26] |
|  | *Theobroma cacao* | JGI | v1.1 | 3641 | [26] |
| Fungal species | *Eremothecium gossypii* | JGI<sup>1</sup> | ATCC10895 | 284811 | [34] |
|  | *Debaromyces hansenii* | JGI | CBS767 | 284592 | [35] [36] |
|  | *Kluyveromyces lactis* | JGI | CLIB210 | 284590 | [35] |
|  | *Saccharomyces cerevisiae* | SGD<sup>2</sup> | S288C | 559292 | [37] |
|  | *Yarrowia lipolytica* | JGI | CLIB122 | 284591 | [35] |

<sup>1</sup>Joint Genome Institute; <sup>2</sup>Saccaromyces Genome Database

OMGene was assessed in three ways: RNA-seq data was used to compare the quality of genes before and after application of OMGene, from both coverage (i.e. the proportion of the predicted gene that is encompassed by reads mapped from RNA-seq data) and splice junction perspectives. To assess the accuracy of start codon prediction, OMGene-modified gene models were subject subcellular localisation prediction and the results were evaluated for consistency across the orthogroup. The RNA-seq data used to assess the success of OMGene were downloaded from the NCBI Sequence Read Archive [23] and are listed in Table 2.

**Table 2:** SRA RNA-seq data sources

|  | Species | SRA ID | Instrument/details | Genes in original annotation |  |  |
| --- | --- | --- | --- | --- | --- | --- |
|  |  |  |  | Total | W/ reads | % |
| Plant species | *A. thaliana* | SRR3932355 | Illumina HiSeq 2500, paired end. Wild type Columbia rep1 | 27416 | 26110 | 95.2 |
|  | *B. rapa* | SRR2984945 | Illumina HiSeq 2000, paired end. ga-deficient dwarf (gad1-2)<br>+GA rep2 | 40492 | 35793 | 88.4 |
|  | *C. papaya* | SRR3509576 | Illumina HiSeq 2500, paired end. SunUp/Sunset cultivar,<br>young hermaphrodite leaf | 27751 | 24589 | 88.6 |
|  | *C. rubella* | SRR797557 | Illumina Genome Analyzer IIx, paired end | 26521 | 21239 | 80.1 |
|  | *T. cacao* | SRR3217315 | Illumina HiSeq 2000, paired end. Flower/leaf sample | 29452 | 25758 | 87.5 |
| Fungal species | *E. gossypii* | N/A<sup>1</sup> | N/A | 4768 | N/A | N/A |
|  | *D. hansenii* | SRR1296968 | Illumina HiSeq 2000, paired end | 5781 | 6272 | 92.2% |
|  | *K. lactis* | SRR1200528 | Illumina Genome Analyzer II, single | 5075 | 5076 | 100% |
|  | *S. cerevisiae* | SRR539284 | Illumina HiSeq 2000, paired end | 6560 | 6572 | 99.8% |
|  | *Y. lipolytica* | SRR868669 | Illumina HiSeq 2000, single | 6432 | 6447 | 99.8% |

### Application of OMGene to publicly available datasets

#### Quantities and nature of changes made

The full plant data set produced 3694 SCU orthogroups, containing 18470 genes. Application of OMGene to this test set resulted in gene model changes to one or more genes in 1543 (41.8%) of these orthogroups. In total, 2017 of the inputted genes (10.9%) were altered. Of these altered versions, 154 genes (7.6% of 2017) were present in the original annotation as alternative (non-primary) transcripts for the inputted gene. Figure 2 shows examples of various types of gene model alteration for genes in *A.* thaliana. A full breakdown of per-species change quantities can be found in Table 3, Figure 3 and Figure 4; Table 4 and Figure 5 show the distribution of the types of changes made. All gene models that were changed by OMGene are included in the supplementary material as a set of GTF files.

**Figure 2:**
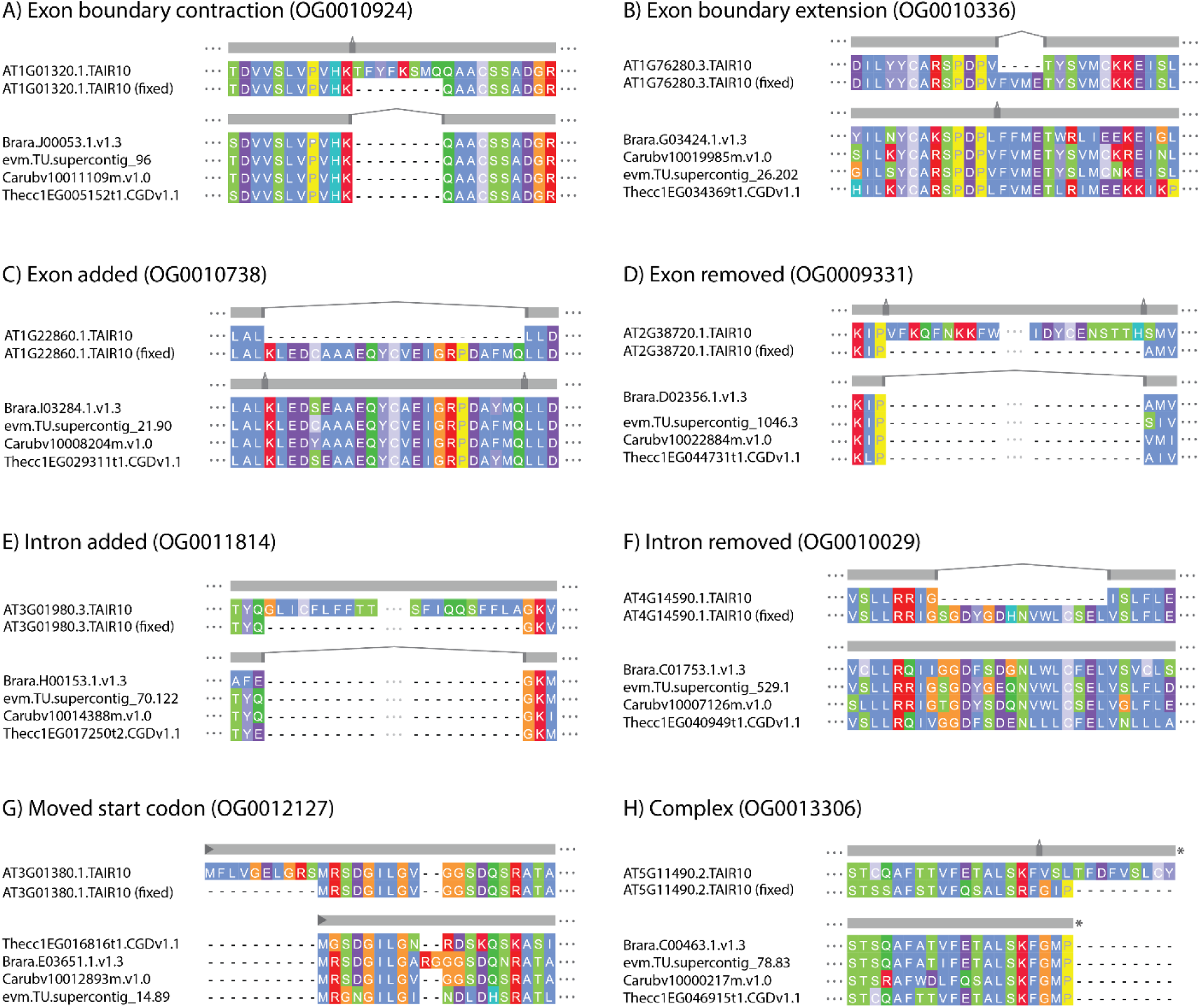
Gene model change examples from *A. thaliana*. Examples of individual gene model changes for genes in *A. thaliana*. A) AT1G01320.1.TAIR10, orthogroup OG0010924, exon extension, splice acceptor side; B) AT1G76280.3.TAIR10, orthogroup OG10336, exon contraction, splice acceptor side; C) AT1G22860.1.TAIR10, orthogroup OG0010738, novel exon introduced; D) AT2G38720.1.TAIR10, orthogroup OG0009331, removed exon; E) AT3G01980.3.TAIR10, orthogroup OG0011814, novel intron introduced; F) AT4G14590.1.TAIR10, orthogroup OG0010029, intron removed; G) AT3G01380.1.TAIR10, orthogroup OG0012127, moved start codon; G) AT5G11490.2.TAIR10, orthogroup OG0013306, complex event: exon has been removed and the previous exon boundary has been extended to include the stop codon.

**Figure 3:**
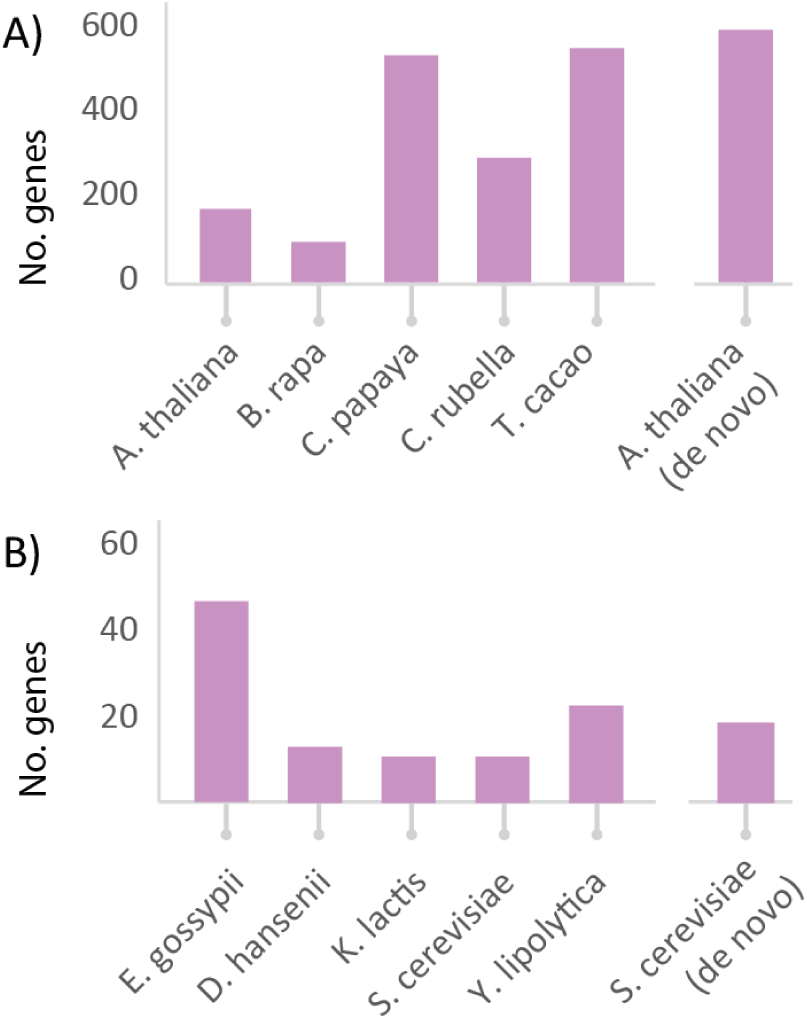
Number of changed genes per species. Chart showing the number of changes made. A) *C. papaya* and *T. cacao* experienced the most changes in the plant data set. The *de novo* version of the *A. thaliana* genome underwent three times more changes than the publicly available one. B) The number of changes made was significantly less for the fungi data set. As in the plants, the representative species *S. cerevisiae* underwent more changes than the public version.

**Figure 4:**
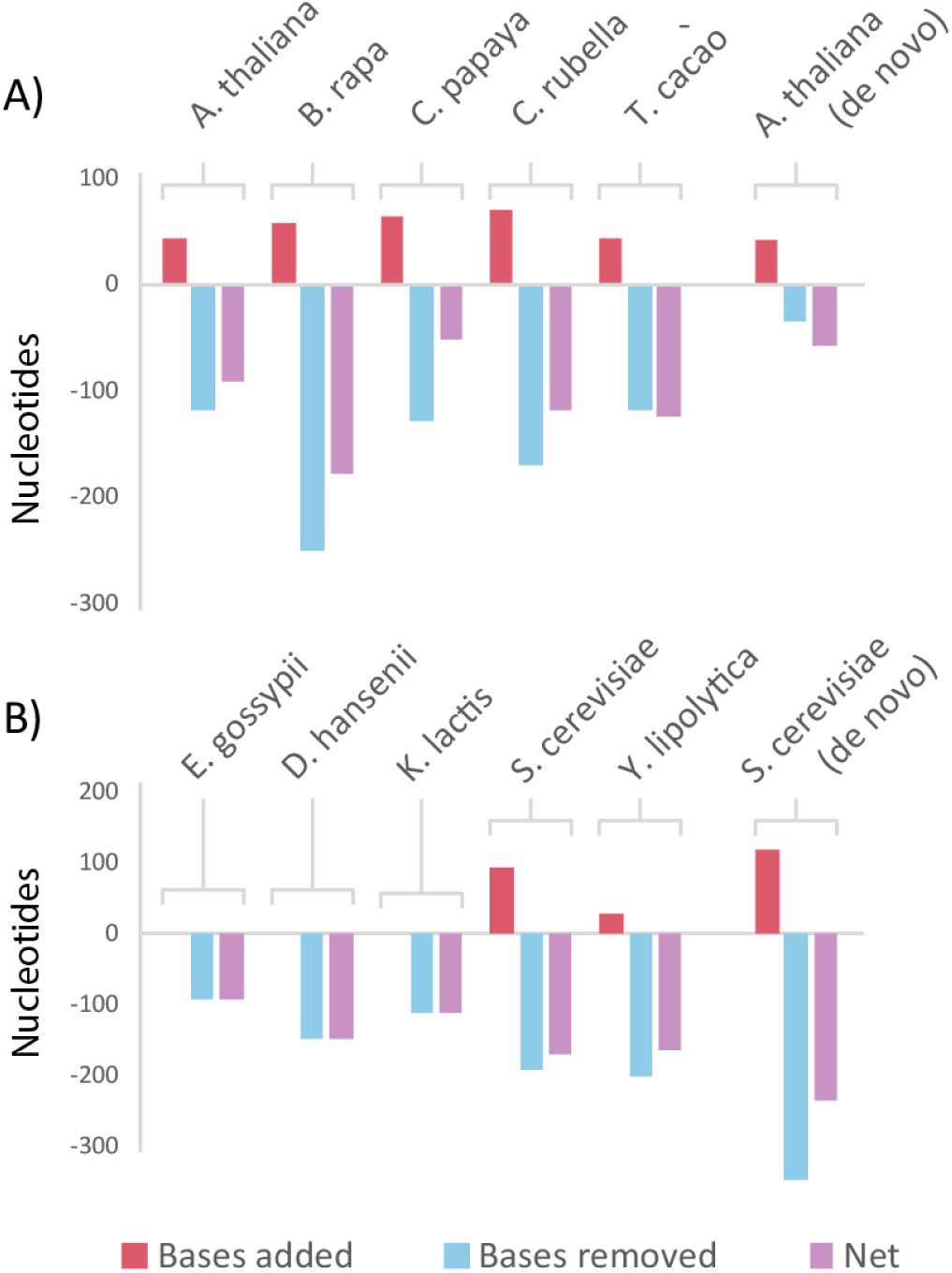
Mean magnitude of changes made. A) Average magnitudes of each change for plants. B) Average magnitudes for changes made to fungal genes.

**Figure 5:**
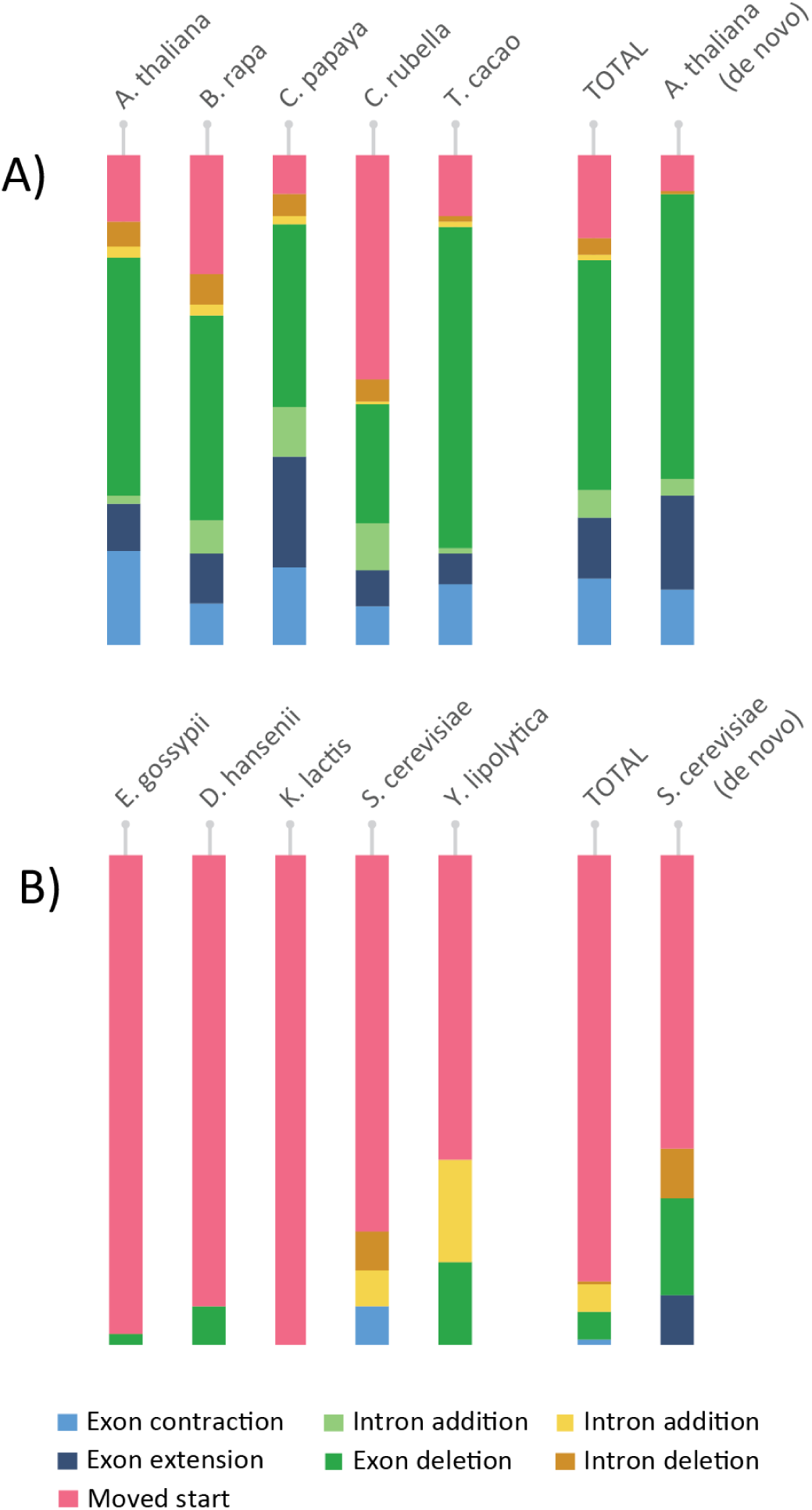
Change type distributions for plant and funal genes. Distribution of types of changes made in the two data sets. A) The most common change in plants was exon deletion. B) In fungi, the most common change was overwhelmingly a moved start codon.

**Table 3:** Per-species gene change breakdown

|  | Species | No. changed genes | Nucleotides added/removed (means per change) |  |  | In original annotation as alternative “non-primary” gene model |
| --- | --- | --- | --- | --- | --- | --- |
|  |  |  | + (mean) | - (mean) | Net (mean) |  |
| Plant species | *A. thaliana* | 175 | 1749 (42.7) | -23747 (-118) | -22139 (-92) | 53 |
|  | *B. rapa* | 97 | 1787 (58) | -25740 (-250) | -23953 (-179) | 4 |
|  | *C. papaya* | 540 | 23820 (65) | -72053 (-128) | -48233 (-52) | 0 |
|  | *C. rubella* | 298 | 6568 (71) | -55005 (-170) | -48437 (-117) | 2 |
|  | *T. cacao* | 556 | 3700 (43) | -120984 (-118) | -117284 (-124) | 95 |
|  | **TOTAL** | 1666 | 37624 (61) | -297529 (-145) | -259905 (-97) | 154 |
|  | *A. thaliana de novo* | 598 | 13623 (42) | -167038 (-35) | -51177 (-57) | N/A |
| Fungal species | *E. gossypii* | 46 | 0 (0) | -4338 (-93) | -4338 (-93) | N/A |
|  | *D. hansenii* | 13 | 0 (0) | -2080 (-149) | -2080 (-149) | N/A |
|  | *K. lactis* | 11 | 0 (0) | -1314 (-110) | -1314 (-110) | N/A |
|  | *S. cerevisiae* | 11 | 93 (93) | -2483 (-191) | -2390 (-170) | N/A |
|  | *Y. lipolytica* | 23 | 117 (29) | -4186 (-199) | -4069 (-163) | N/A |
|  | **TOTAL** | 104 | 210 (42) | -14401 (-135) | -14191 (-127) | N/A |
|  | *S. cerevisiae de novo* | 19 | 601 (120) | -5561 (-347) | -4960 (-236) | N/A |

**Table 4:**
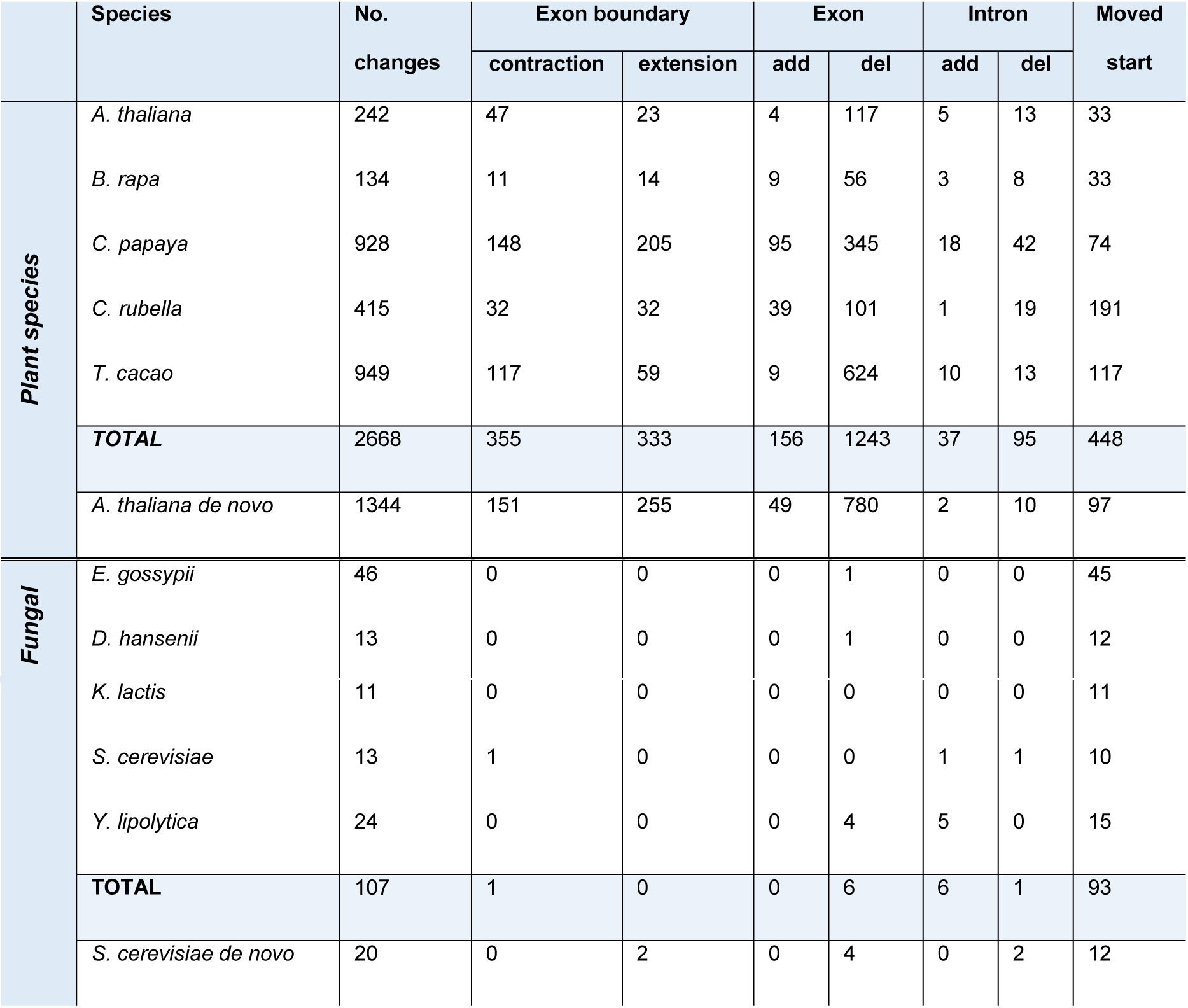
Summary of gene model change categories

The plant species that experienced the highest number of changes were *C. papaya* and *T. cacao*, which is consistent with them being more recently published and less well-studied genomes. For all species, more nucleotides were removed than were added, indicating either that gene models predictions tend to be over-cautious or that OMGene is more proficient at removing material than at adding it in. In terms of the types of changes made, exon deletion was by far the most commonly seen change, followed by moved start codon and exon boundary adjustment (Figure 5). It should be noted that exon deletion events also encapsulate the separation of erroneously fused gene models, which can contribute many exon deletion events simultaneously.

For the full fungal data set, 2710 SCU orthogroups were considered, containing 13550 genes. Of these, 100 orthogroups (3.7%) exhibited some change, and 109 genes (0.8%) were altered. As above, a full breakdown of per-species change quantities can be found in Table 3, Figure 3, and Figure 4 with the full distribution of change types shown in Table 4 and Figure 5. In this case, *E. gossypii* was the most commonly altered proteome, consistent again with it being one of the lesser-studied species on the list. By far the most common change type in the fungal data set was a moved start codon, consistent with the fact that splicing is a rare event in fungal genes (on average 5.09 exons for plants, 1.08 exons for fungi).

To simulate a *de novo* genome annotation project, OMGene was also applied to plant and fungal data sets with *de novo* predicted gene models for representative species, *A. thaliana* and *S. cerevisiae*. These species were chosen as they have the most complete annotations of their respective data sets, and therefore these genes are likely to be the most reliable for training a gene finding algorithm. The genome annotation tool used was Augustus (see Methods) as it is one of the best and most frequently used gene prediction algorithms.

For the plants data set with Augustus predictions for *A. thaliana*, 3694 SCU orthogroups were considered. Of these, 598 (16.2%) saw some change in an *A. thaliana* gene. For the fungi data set, 2710 SCU orthogroups were considered. Of these, 19 (0.7%) saw some change in a *S. cerevisiae* gene. Table 3 and Table 4 show a full breakdown of the types and amounts of changes made. As expected, in both cases, the total number changes and the average size of change made is greater for the *de novo* predicted gene models than the curated gene models. However, the distribution of types of changes made remained roughly the same.

#### Splice junction and feature coverage analysis

To assess the validity of changes made by OMGene, both the original and the updated gene model sets were compared using publicly available RNA-seq data from the NCBI Sequence Read Archive [23] (see Methods and Table 2). Each amended gene was assessed in two ways relative to this data: firstly by comparing the exact splice junction locations with RNA-seq derived splice junctions; secondly by evaluating the coverage of exonic regions with RNA-seq. To control for unreliable data, some genic regions were omitted from this analysis. Gene regions in which the RNA-seq data suggested there were indels in the reference genome, or that were within 1000bp of the end of a contig or scaffold, or that contained 10 or more contiguous “N” nucleotide bases were omitted from the analysis (see Methods). Regions with these characteristics prevent the creation of reliable gene models, and so are not useful for determining gene model accuracy.

Gene models outputted by OMGene were assessed on whether or not their junction and coverage F-scores (see Methods) had improved or been reduced. The full results can be seen in Table 5. For the plant data set, OMGene improved the agreement of the gene model with the splice junctions inferred from RNA-seq data for 729 genes, while 125 gene models exhibited reduced agreement (85.3% improved). Similarly, when assessing RNA-seq coverage of gene models OMGene improved the agreement of the models with the data for 1026 genes, while 167 genes exhibited reduced agreement (86.0% improved). For the *de novo* predicted *A. thaliana* genes, the success rates were essentially the same as for the public data (87.3% and 91.1% improved by junction and coverage F-scores respectively), but the absolute quantity of genes exhibiting a changed score increased roughly four-fold. This difference represents the considerable effort and evidence-based curation that has been invested in the *A. thaliana* genome annotation.

**Table 5:** RNA-seq coverage and junction F-scores

|  | Species | Junction F-score |  | Coverage F-score |  |
| --- | --- | --- | --- | --- | --- |
|  |  | Better | Worse | Better | Worse |
| Plant species | *A. thaliana* | 94 (87.8%) | 13 (12.1%) | 109 (91.5%) | 10 (8.4%) |
|  | *B. rapa* | 24 (63.1%) | 14 (36.8%) | 29 (56.8%) | 22 (43.1%) |
|  | *C. papaya* | 246 (82.2%) | 53 (17.7%) | 344 (83.9%) | 66 (16.0%) |
|  | *C. rubella* | 90 (89.1%) | 11 (10.8%) | 186 (91.6%) | 17 (8.3%) |
|  | *T. cacao* | 275 (88.9%) | 34 (11.0%) | 358 (87.3%) | 52 (12.6%) |
|  | **TOTAL** | 729 (85.3%) | 125 (14.6%) | 1026 (86.0%) | 167 (13.9%) |
|  | *A. thaliana de novo* | 422 (87.3%) | 61 (12.6%) | 475 (91.1%) | 46 (8.8%) |
| Fungal species | *D. hansenii* | 1 (100.0%) | 0 (0%) | 4 (66.6%) | 2 (33.3%) |
|  | *K. lactis* | 0 (N/A) | 0 (N/A) | 9 (100.0%) | 0 (0%) |
|  | *S. cerevisiae* | 0 (N/A) | 0 (N/A) | 6 (75.0%) | 2 (25.0%) |
|  | *Y. lipolytica* | 2 (28.5%) | 5 (71.4%) | 11 (64.7%) | 6 (35.2%) |
|  | **TOTAL** | 3 (37.5%) | 5 (62.5%) | 30 (75.0%) | 10 (25.0%) |
|  | *S. cerevisiae de novo* | 4 (100%) | 0 (0%) | 11 (64.7%) | 6 (35.2%) |

The results for the fungal data set (see Table 6) were not as good. Notably very few gene models showed any change in junction F-score, with only 8 genes exhibiting a changed score. This is due to the relatively simple exon structure of fungal genes, for which splicing is very rare, and splicing events predicted by OMGene are much less likely to be correct. In this case 3 genes had an improved score, and 5 had a reduced score (37.5% success), with all 5 of the losing genes coming from *Y. lipolytica*. The most common change made to fungal genes was a moved start codon, which, although not detectable in the junction F-score, can be detectable in the coverage F-score. This is reflected in the results, where 30 genes showed an improved coverage F-score and 10 genes showed a worse coverage F-score (75% improved). In the *de novo* case, again the numbers increased while the percentage success remained roughly the same, with 4 (100%) genes improving by junction for *S. cerevisiae* and 11 (64.7%) improving by coverage score. The highly compact nature of fungal genomes, with few exons and limited space between genes means that the accuracy of *de novo* predicted genes is higher than in plants. Thus the utility of OMGene on these comparatively simpler genomes is limited.

**Table 6:**
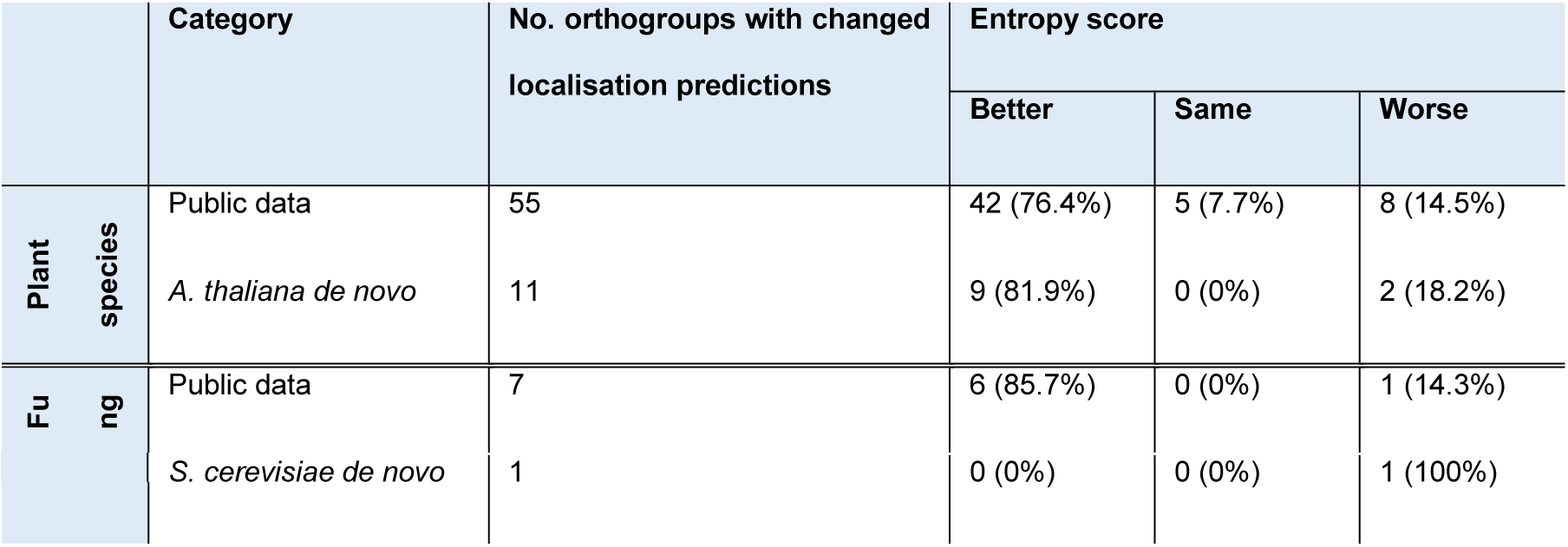
Subcellular localisation predictions.

|  | Category | No. orthogroups with changed localisation predictions | Entropy score |  |  |
| --- | --- | --- | --- | --- | --- |
|  |  |  | Better | Same | Worse |
| Plant species | Public data | 55 | 42 (76.4%) | 5 (7.7%) | 8 (14.5%) |
|  | *A. thaliana de novo* | 11 | 9 (81.9%) | 0 (0%) | 2 (18.2%) |
| Fungal | Public data | 7 | 6 (85.7%) | 0 (0%) | 1 (14.3%) |
|  | *S. cerevisiae de novo* | 1 | 0 (0%) | 0 (0%) | 1 (100%) |

Many of the cases for which OMGene results differ from RNA-seq evidence are attributable to real biological variability that confounds the evaluation criteria of the algorithm. For example, there are some instances where the most evolutionary conserved splice site was not the splice site observed in the RNA-seq data. Such events, by definition, cannot be detected by OMGene. Furthermore, RNA-seq mapping errors also contributed to reduced scores, as did artefacts resulting from spliced UTRs, and jagged read profiles, particularly in the fungal data, that made some coverage scores difficult to calculate reliably. Finally, the presence of multiple transcript isoforms within the RNA-seq data can reduce the score for a valid transcript even if it is the best choice for that particular gene. While users of OMGene should be aware of these confounding factors, the above data demonstrates that, in general, OMGene is much more likely to improve a given gene model than not.

#### Assessment of subcellular localisation predictions for 5’ end analysis

Given that genes from the same orthogroup are, by definition, assumed to be evolutionarily related, it is reasonable to assume that they should be consistent in their predicted subcellular localisation. Several sub-cellular targeting sequences are located at the N-termini of genes [24], thus one expects genes with inaccurately predicted start codons to yield inaccurate results when assessing their targeting signals. Genes belonging to orthogroups changed by OMGene were assessed to determine whether the changes resulted in increased consistency of their predicted subcellular localisation characteristics of all genes in the orthogroup. Targeting predictions were made using TargetP [25], and Shannon entropy was calculated to assess the consistency of the predictions within the orthogroups (see Methods). Entropy scores were compared only for orthogroups in which at least one gene model was altered by OMGene. An entropy score of 0 indicates that all members of the orthogroup are predicted to localise to the same sub-cellular compartment; the worst possible entropy score given five genes and four possible localisations identified by TargetP (chloroplast, mitochondrion, secreted, cytoplasmic) is 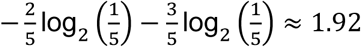, indicating that only two of the genes agree. An example orthogroup whose prediction entropy score has been improved by start codon adjustment can be seen in Figure 6**Error! Reference source not found..**

**Figure 6:**
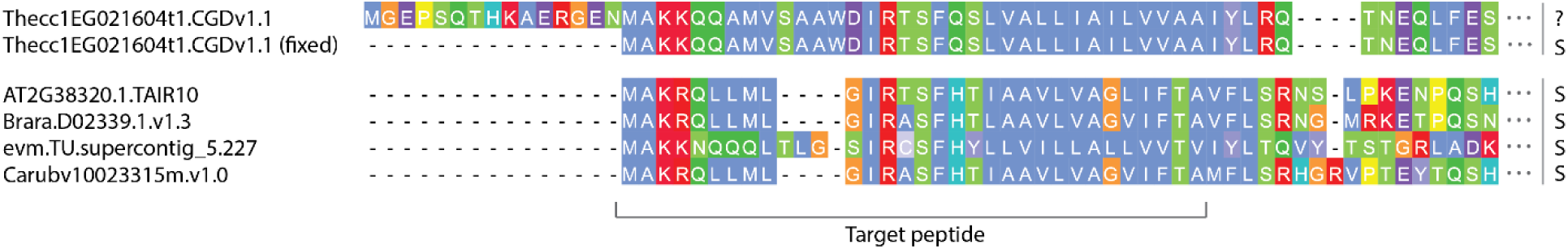
Example change in subcellular localisation prediction. Example change in subcellular localisation prediction for a gene. Thecc1EG021604t1.CGDv1.1 from *T. cacao* has undergone a change in start codon, revealing a signalling peptide at its 5’ end. In this case, what was previously assumed to be cytosolic has been found to target the secretory pathway, the same as the other members of the orthogroup (OG0009265). In this case, the Shannon entropy score for the orthogroup has fallen from 0.72 to 0.

The 1543 plant orthogroups in which one or more genes were altered were subject to subcellular prediction analysis. Of these, gene model changes made by OMGene resulted in changes in predicted subcellular localisation for one or more constituent members of 55 orthogroups. In total, 74 improved agreement between gene models (74%), 13 remained the same (13%), and 13% increased entropy and thus increased disagreement between gene models. In contrast, for the fungal dataset only 7 out of 95 changed orthogroups exhibited a change in subcellular localisation prediction, with 6 of these changes improving the consistency of localisation prediction (85.7%) and 1 increasing disagreement (14.3%). Similar results were obtained for the simulated *de novo* annotation analysis in plants, although again the data were sparse here. Orthogroups containing the *de novo* predicted *A.* thaliana gene were considered together with the four original genes for the other species. Here, 11 of the *A. thaliana* genes experienced a change in subcellular localisation following application of OMGene. Of the 11 orthogroups containing these, 9 improved consistency (81.9%) and 2 reduced the consistency (18.2%). For the fungal data set, the data was extremely sparse, with only one gene experiencing a change in its targeting prediction, which reduced the consistency for its parent orthogroup. Thus, although data were sparse for the fungal dataset, in both the fungi and plant dataset the consistency of gene models was improved from a subcellular targeting perspective.

## Discussion

Here we present OMGene, an automated method for improving the consistency of gene model annotations across species. OMGene is intended for use in computational *de novo* genome annotation projects where no empirical data (such as RNA-seq data) is available to train or correct gene model predictions, or to assist the construction of gene models for genes that are not expressed in the data available. OMGene is also designed to help users who wish to leverage conservation information to correct gene models of a single gene of interest across a set of species. Thus OMGene is suitable for both large and small scale analyses.

### OMGene results reflect differences in gene model complexity between species sets

To demonstrate the utility and performance characteristics of OMGene, it was applied to two separate datasets of well-annotated plant and fungal genomes. When applied to the plant data set, OMGene altered the gene models of one or more genes in 41.8% of the orthogroups that were evaluated. In contrast, only 3.7% of orthogroups were subject to modification in the fungal data set. This result reflects the differences in gene model complexity between the two species groups. Specifically, gene models in plants tend to have more exons than fungi (mean = 5.09 exons for plants, 1.08 exons for fungi) and thus there is considerably more potential for gene model variation in plants than in fungi. In light of this it was unsurprising that the most frequently observed change made in fungi was a change in choice of start codon. This is also reflected in the high number of removed exons from plant genes, which is contributed to partly by the separation of erroneously fused adjacent genes.

### OMGene works well on complex gene models

The changes made by OMGene were assessed relative to splice-mapped RNA-seq data to assess the level to which it had improved the gene models. For the plant data set, the results from OMGene clearly resembled the empirical data much more closely on the whole, with 85.4% and 86.0% of genes improving in terms of their splice junctions and their coverage respectively. The profiles were different for different species, with many more changes being made for *C. papaya* and *T.cacao;* in addition the number of successes for *B. rapa* was slightly lower than for the other species.

The number of junction changes made for the fungal data set was considerably lower: only 8 changed genes had an altered junction F-score, 62.5% of which become worse after OMGene. Though this is less than the plant data set, it should be noted that the resolution of this data set does not lend itself to accurate conclusions about the general validity of changes made to fungal genes. The resolution and success rate for fungal genes from a coverage perspective was slightly higher, with 75% of the genes with changed scores improving. The low resolution of junction data for fungal genes reflects the rarity of complex gene models in these species, and thus the low likelihood that deviations from simple, single-exon gene models are correct. Thus, while OMGene does not always produce gene models that agree optimally with transcriptome data, it does improve the overall quality of gene model annotations even for relatively simple fungal genomes.

The improvements in gene model accuracy made by OMGene for the *de novo* predicted proteomes were much the same as for the publicly available, curated genes models. However, the number of changes made to the *de novo* predicted set was much greater, indicating that the considerable labour that has been applied to these model organisms has successfully controlled for potential errors. It should be noted that, although OMGene managed to improve many of the gene models outputted by Augustus, the two agreed in most cases (86.1% and 98.6% for plants and fungi respectively), indicating that the basic implementation of a well-trained Augustus *de novo* prediction produces genes that are highly consistent with their orthogroups.

### OMGene improves the consistency of subcellular localisation predictions

In addition to assessment of splice junctions, gene models were assessed by the consistency of their predicted subcellular localisation. Given that the orthogroups used in this analysis comprise ubiquitously conserved single copy genes, it is logical to assume that these genes should generally have the same subcellular localisation. For the full plant data set, of all orthogroups whose genes had different subcellular targeting predictions after application of OMGene, 76.4% had improved intra-orthogroup consistency, with 85.5% either improving or remaining the same. For the full fungal data set, although the data were sparse, 85.7% of the orthogroups considered had improved consistency.

The results for the plant data set were similar for the *de novo* annotated set (85.7% improvement). For fungal orthogroups containing *de novo* predicted *S. cerevisiae* genes, the only gene whose localisation prediction changed caused the consistency of its orthogroup to decrease, however the resolution of the data in this case is not sufficient to draw any conclusions. Thus, application of OMGene improves the accuracy of start codon specification in gene models.

### Conclusion

When applied to publicly available plant and fungal data sets, OMGene demonstrates proficiency in improving gene models from multiple perspectives. The overall improvement is larger for genomes with complex gene models.

## Methods

### Algorithm description

The input for OMGene is a set of GTF gene model files and a set of corresponding FASTA genome files. There should be one GTF per FASTA file, and each GTF should contain the coordinate information for a single gene. If the GTF contains multiple transcript variants then these are considered together as variants of a single gene.

For each inputted gene, the algorithm defines its *gene region* to be the region spanning the first and last base of any of its corresponding gene models, with a user-selected number of buffer bases either side (default value is 600bp). The initial step of OMGene is to cross-align the amino acid sequences from each gene with the gene regions of the other genes, using Exonerate [21]. The rationale behind this step is to find exonic regions that are present in one or more gene models but absent from one or more annotated gene model. This is performed three times: first by cross-aligning the input protein sequences against all gene regions, second by cross aligning the protein sequences that have been found in the first step against all gene regions, and finally by cross aligning all individual exon sequences from the first step. This three-step process mitigates against lack of detection due to gene model errors in one or more of the input genes. This, together with the exons from the original gene sequences, comprises a set of potential gene parts, which may overlap and which may be incompatible in reading frame. Compatible combinations of gene parts (i.e. without frame-shift errors) are strung together to form a putative gene model. Many such putative gene models may exist: the set of putative gene models with the highest alignment score (see alignment score calculation below) is carried forward to the next step.

The set of putative gene models from the previous step are aligned, and the set of putative exons from all genes is divided into *adjacency groups*: sets of exons that overlap each other in the alignment (see below). Exons are added in sequentially in these adjacency groups, and at each stage a valid gene model is sought on the left hand side of the gene (i.e. starting at the start codon and seeking to adjoin exons in valid donor-acceptor pairs). Multiple options for each gene are produced at each new junction, by recursively seeking out, or “wiggling” splice junctions (or start codons) in each frame either side of the existing exons start and end points. This produces a set of junction options for each pair of exon ends. A multipartite choice function is then used to choose the best option for each pair of exons, as described below. In the event that a particular exon is very small (<40bp), or does not yield any valid junction sites, both that exon and the one before it are probed for removal, and the variant with the removed exon is compared against the other partial gene models in the evaluation step. Once this recursive step ceases to produce new gene modes, the gene model set with the highest alignment score is declared the winner, and the next putative exon from the next adjacency group is added. This is repeated until there are no further exons to add.

To ensure that the optimisation process did not overlook potentially better variants in the user-supplied gene models, the process above is repeated. This time, instead of varying exons start and end sites, the set of newly created junctions are compared against the original junctions, aiming to find the optimal combination of new and old junctions.

The final step involves filtering the changes based on a selection of categories that have been observed to over-fix gene models. Firstly, we require the alignment score *α* of a 10 amino acid region each side of the change to have either remained the same or improved. This is a basic requirement which should be met in most cases due to the way in which sequence variants are chosen. Secondly, changes that have opened gaps in the alignment of three or more of the sequences are not allowed: this is a common occurrence due to sequences proximal to exon termini that that by chance feature valid splice junction sequences that are in frame with the adjacent exons and are evolutionarily conserved. These tend not to be correct. Thirdly, very small changes are forbidden: changes that have resulted in two or fewer amino acids being changed in a gapless region of the alignment, such that the new alignment is also gapless, are ignored. Similar changes to larger regions require an *α* increase of 4 or more. This is to avoid changes that reflect multiple choices of donor-acceptor pairs for essentially identical sequences. Thirdly, the alignment in the region of the change must be of reasonable quality: for unchanged 5 amino acid regions near the change, the adjusted alignment score 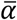 must be 3 or higher (or all gaps) for some subset of three sequences containing the sequence of interest. Similarly the resulting score for the changed region must also be higher than 3 or all gaps. Exon boundaries that do not pass the filters are discarded and the genes are reconstructed a final time, allowing only the surviving boundaries and those that were present in the original gene. The resultant genes are outputted in GTF, amino acid FASTA and CDS FASTA format.

### Data sources

For algorithm development and evaluation, a set of five small, well-annotated fungal genomes and a set of five well-annotated plant genomes (Table 1) were selected. Orthogroups were inferred using OrthoFinder [20]. For the plant data set, where multiple transcript variants were available, the primary transcript was used as listed in Phytozome [26]. RNA-seq data sources are listed in Table 2, and were downloaded from the Sequence Read Archive [23].

### *De novo* gene prediction

*De novo* gene predictions were made using Augustus [22] version 3.2.2. Training was performed using all well-formed gene models from each species, and using the autoAugTrain.pl script included with the software. Augustus was run individually on each genome with the default settings.

### Alignment score

An amino acid alignment can be considered as an ordered sequence 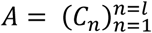 of columns 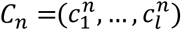. The *column score γ* for a column *C*_*n*_ is defined as the average pairwise Blosum62 score for amino acids in that column:

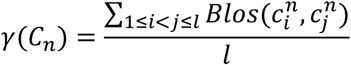

The Blosum62 matrix was used as it is the basis for the MAFFT alignment algorithm. The *alignment score α* for an alignment *A* is constructed column-wise as:

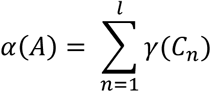

The *adjusted alignment* score 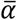 is defined as 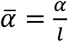, where *l* is the alignment length.

### Multipartite choice function

The multipartite choice function (Figure 7) aims, for a set of *k* gene regions and a set of *l*_*k*_ gene model variants for each gene region, to choose an optimal set containing one gene model variant from each gene region such that the alignment score is maximised. This problem is equivalent to finding the heaviest maximal clique in an edge-weighted complete multipartite graph.

**Figure 7:**
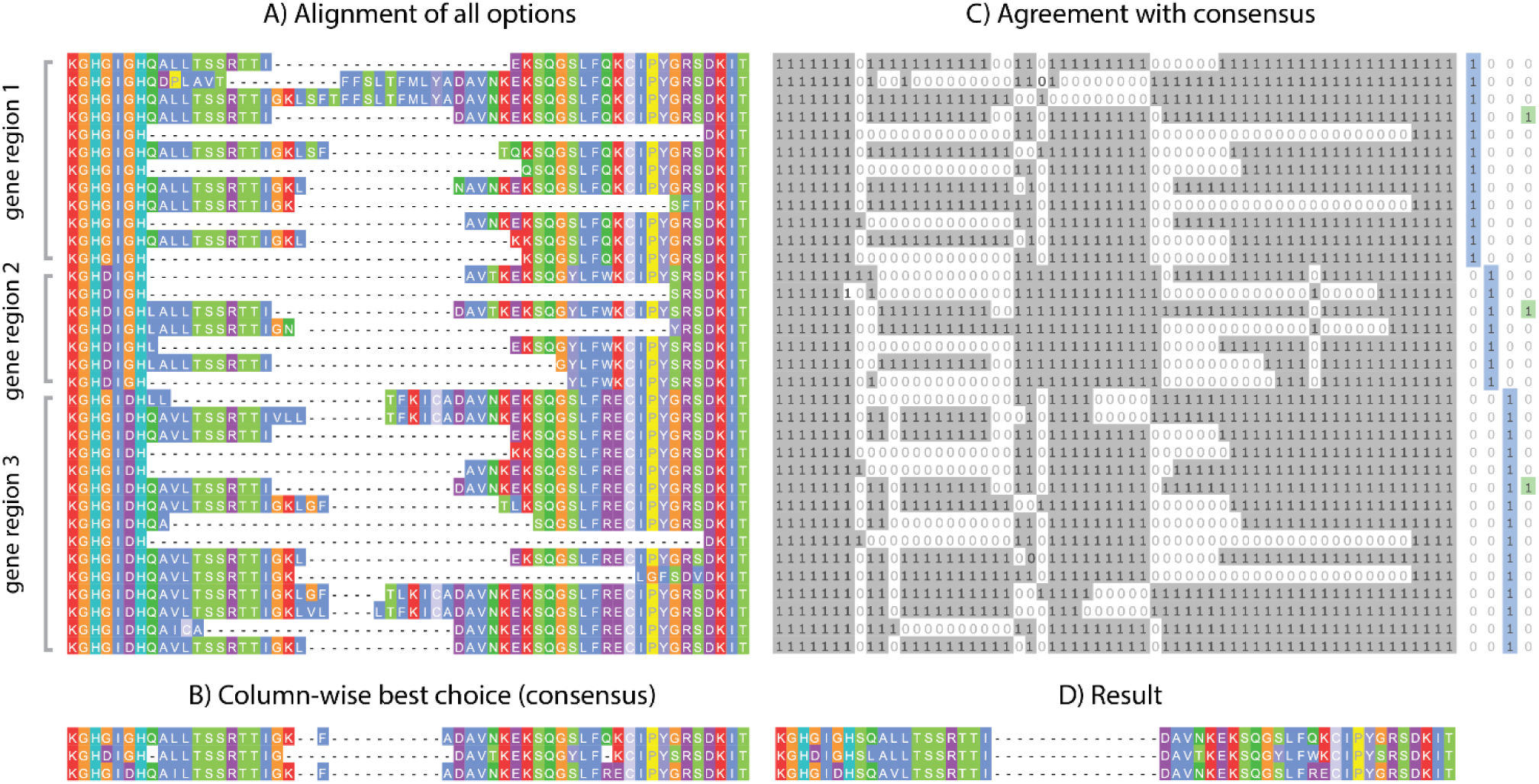
Multipartite Choice Function. The choice function aims to find optimal variants from a set of protein sequences. A) Sequences are aligned; B) A consensus alignment is produced: on a column-by-column basis the choice of amino acid for each sequence that optimises the alignment score for that column is chosen as a representative; C) A binary representation is produced from the original alignment: for each base in alignment, a 1 is assigned if the base matches the consensus, and a 0 is assigned if it does not. This leaves a sequence of vertical binary strings. The aim is to find a single vertical binary string that agrees with (i.e. is a bitwise subset of) as many as possible of these, and that is also compatible with the category constraints. The best such string in this case is shown to the right in green. D) The result.

To reduce the complexity of the problem, options are chosen by comparison with a reference consensus alignment, produced by taking the most consistent set of amino acids for each column in a global alignment individually (Figure 7A-B). This column-wise optimisation is fast, and provides a basis for the sequence-wide optimisation. To produce the consensus, The set of Σ*l*_*k*_ options is aligned to the reference (the original alignment) using MAFFT -add [27]. The inconsistent regions are then isolated and re-aligned using the more accurate but more computationally intensive MAFFT l-ins-i. For each column in the alignment, the set of amino acid choices (one for each gene region) that optimises the alignment score for that column is chosen as the consensus.

For each option *i* a binary string 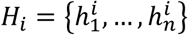 is produced describing for each position in the alignment whether or not that option matches the consensus (Figure 7C). The chosen subset will be the set of options that globally maximises agreement with the consensus. If the strings {*H*_*i*_}_*i*_ are stacked vertically, such that they can be read as columns 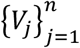 then the task is equivalent to finding a columnar binary string *V* with one nonzero entry for each gene region such that |*V*_*i*_:*V*⊆*V*_*i*_| is maximised.

Given the set 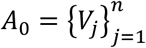, an optimal subset is deduced by sequential random sampling. Ignoring all-1 strings, an initial *W*_0_ = *V*_*k*_ is chosen at random from *A*_0_. For sets *S*_1_, *S*_2_ and a set of “checkpoints” *R*, the set *S*_1_ is *compatible with S*_1_ respect to *R* = {*R*_*i*_}_*i*_ if the binary intersection *S*_1_ ∩ *S*_2_ ∩ *R*_*i*_ is nonzero for all *i*. Define *A*_*n*_ = {*a* ∩ *W*_*n*−1_:*a*,*W*_*n*−1_ compatible w.r.t *G*}, where *G* is the set of binary strings which are zero for all but one gene region, at each stage choosing *W*_*n*_ at random from *A*_*n*_. The process *A*_0_, *A*_1_, *A*_2_, … eventually converges on a single binary string. This reduction is performed a user-selected number of times, the default being 1000. The result that is a subset of the largest number of *V*_*i*_ is declared the winner. In the event that the result still contains more than one option for each gene region, subsets of options are calculated and their multiple alignment score *α* is calculated, the winner being the subset with the highest *α*. In the event that multiple subsets exhibit the same maximal *α*, a subset is chosen arbitrarily from them.

### Adjacency group calculation

OMGene builds genes sequentially by iteratively adding in putative exons to multiple genes simultaneously. Care must be taken to ensure the gene parts (which in turn become exons once gene models are constructed) are added in a way conducive to vertical comparison of relevant regions (see Figure 8). In OMGene, gene parts are considered in sequential *adjacency groups* based on their coordinates in a multiple sequence alignment. Prototype gene models are formed by stringing together amino acid sequences for individual putative exons for each gene region: these are then aligned, and a graph is formed from this alignment. Each putative exon is a node on the graph, and two exons are connected by an edge if one of the exons overlaps the other by a third or more of its length. The adjacency groups are then defined to be cliques in this graph. Cliques are determined using the python implementation of the NetworkX package [28].

**Figure 8:**
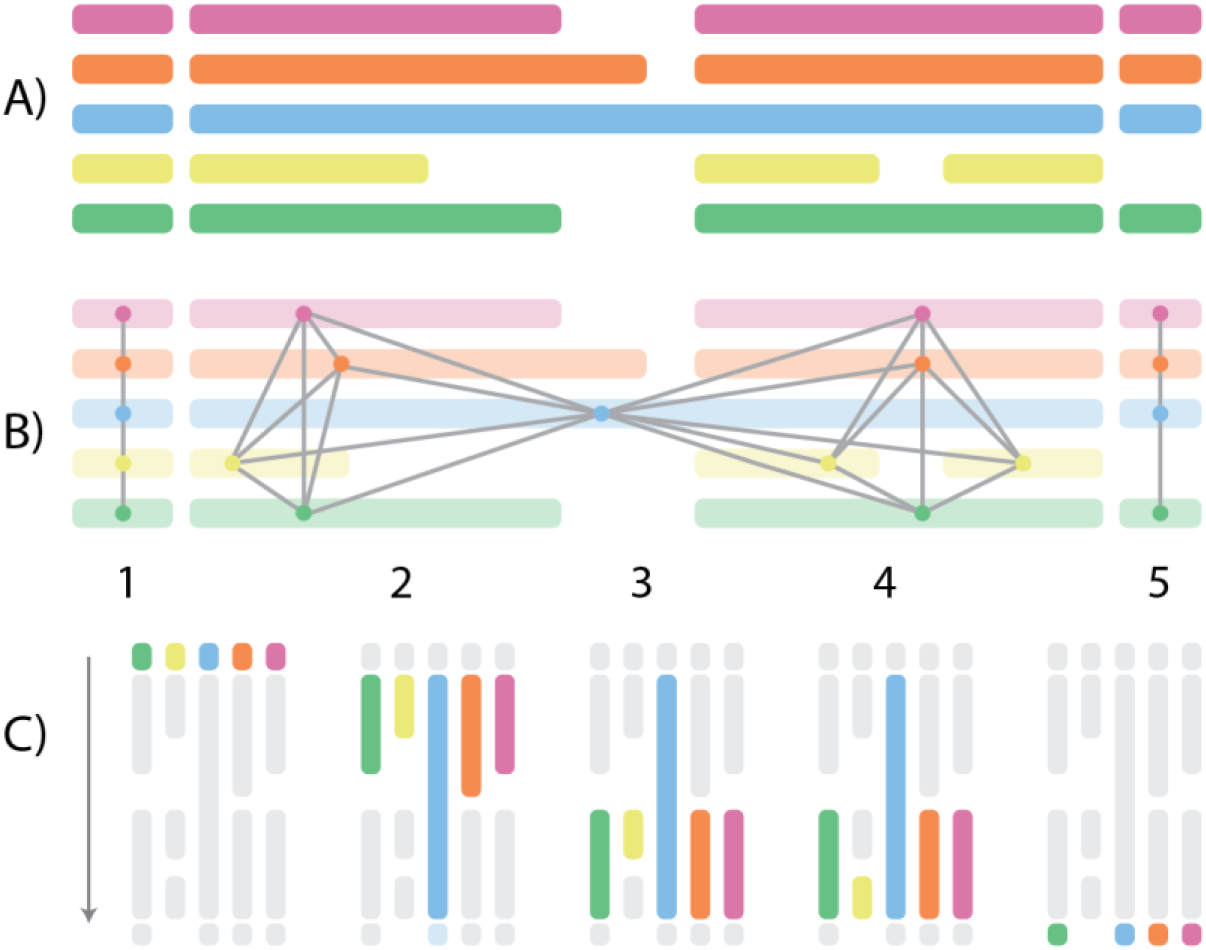
Adjacency group calculation. Calculation of adjacency groups. A) Amino acid sequences for individual putative exons are strung together and aligned. B) A graph is formed with vertices formed by gene parts (or exons), and edges drawn when the overlap between two parts is greater than or equal to two thirds the length of one of them. C) Cliques are extracted and then ordered lexicographically to form the adjacency groups.

### Junction F-score

The *junction F-score* for a gene is a measure of how well the splice junctions observed in mapped RNA-seq data are represented in the gene model. For a gene model *G* and corresponding gene region *R*, define *J*_*G*_ to be the set of individual intron beginning and end coordinates in the gene model, and define *J*_*R*_ to be the set of map junction beginning and end coordinates in the mapped RNA-seq data. A minimum of 10 reads is required for a given RNA-seq junction to be counted. We may then define the junction F-score as:

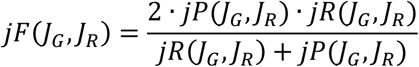

where

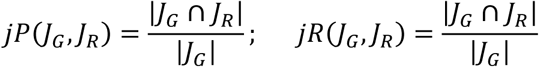

The direction of each junction site (start or end of a junction) is taken into account when considering the intersection of the two sets.

### Coverage score

The *coverage score* is a measure of how well RNA-seq data represents a given gene. Given that gene expression levels can vary considerably and irregularly across the length of a transcript [13]– [16], care must be taken to ensure the expression profile for a gene region is properly interpreted. For example, sample preparation methods can bias coverage towards the centre and 3’ ends of the transcript; furthermore, jagged read profiles and transcription of antisense regions [29] and other intronic ncRNAs can cause expression profiles to be highly non-binary. To mitigate this, a rolling threshold approach is used. For a gene region *R*, and a genomic coordinate *x* ∈ *R*, the expression characteristic *χ* is defined as:

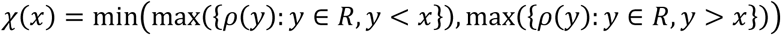

Where *ρ*(*y*) is the read count at genomic coordinate *y*. Bases in the gene region to which the RNA-seq data has been mapped are categorised based on whether they are likely to correspond to exonic or non-exonic regions: a base *x* is considered to be *on* (i.e. likely included in the mature mRNA) if 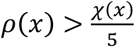, and *off* (i.e. likely not included in the mature mRNA) if 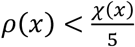. The coverage score for a gene model *G* = {*G*_1_,…,*G*_*n*_} where the *G*_*i*_ are alternately exons and introns, is defined to be:

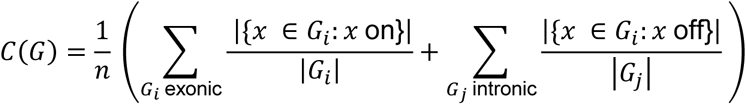

that is, the average length-adjusted coverage score for each individual feature in the gene.

### RNA-seq data

RNA-seq data were downloaded from the Sequence Read Archive, and aligned to the genome with Hi-SAT2 [31], [32] using default parameters. Per-base coverage was calculated using SAMtools mpileup [33].

### Subcellular localisation analysis

Subcellular localisation for both the plant and fungal datasets was determined using TargetP [25]. For the plant dataset only, TargetP was run with the -P option to predict chloroplast targeting sequences. The localistion consistency for an orthogroup *O* was calculated as an entropy score across the categories for each gene:

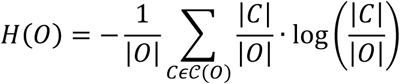

where 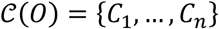 is the partition of genes in *O* into their localisation categories.

## Availability of data and materials

The software is available under the GPLv3 licence at https://github.com/mpdunne/omgene.

## Competing Interests

The authors declare that they have no competing interests.

## Funding

SK is a Royal Society University Research Fellow. This work was supported by the European Union’s Horizon 2020 research and innovation programme under grant agreement number 637765. MPD is supported by an EPSRC studentship through EP/G03706X/1.

## Author’s Contributions

SK conceived the project. MPD developed the algorithm. SK and MPD analysed the data and wrote the manuscript. Both authors read and approved the final manuscript.

